# Chromatin accessibility combined with enhancer clusters activation mediates heterogeneous response to dexamethasone in myeloma cells

**DOI:** 10.1101/2021.09.06.459068

**Authors:** Victor Gaborit, Jonathan Cruard, Catherine Guerin-Charbonnel, Jennifer Derrien, Jean-Baptiste Alberge, Elise Douillard, Nathalie Roi, Magali Devic, Loïc Campion, Frank Westermann, Phillipe Moreau, Carl Herrmann, Jérémie Bourdon, Florence Magrangeas, Stéphane Minvielle

## Abstract

Glucocorticoids (GC) effects occur through binding to the GC receptor (GR) which, once translocated to the nucleus, binds to GC response elements (GREs) to activate or repress target genes. Among GCs, dexamethasone (Dex) is widely used in treatment of multiple myeloma (MM), mainly in combination regimens. However, despite a definite benefit, all patients relapse. Moreover, while GC efficacy can be largely attributed to lymphocyte-specific apoptosis, its molecular basis remains elusive.

To determine the functional role of GR binding in myeloma cells, we generated bulk and single cell multi-omic data and high-resolution contact maps of active enhancers and target genes. We show that a minority (6%) of GR binding sites are associated with enhancer activity gains and increased interaction loops. We find that enhancers contribute to regulate gene activity through combinatorial assembly of large stretches of enhancers and/or enhancer cliques. Furthermore, one enhancer, proximal to GR-responsive genes, is predominantly associated with increased chromatin accessibility and higher H3K27ac occupancy. Finally, we show that Dex exposure leads to co-accessibility changes between predominant enhancer and other regulatory regions of the interaction network. Notably, these epigenomic changes are associated with cell-to-cell transcriptional heterogeneity. As consequences, *BIM* critical for GR-induced apoptosis and *CXCR4* protective from chemotherapy-induced apoptosis are rather upregulated in different cells.

In summary, our work provides new insights into the molecular mechanisms involved in Dex escape.

## Introduction

Dexamethasone (Dex), a synthetic glucocorticoid known for its anti-inflammatory and immunosuppressive activities, in combination with immunomodulatory drugs (IMiDs) and proteasome inhibitors (PIs), is the standard induction treatment in transplant-eligible patients with newly diagnosed multiple myeloma. Recently, the use of a new class of drugs, the monoclonal antibody daratumumab, in combination with thalidomide (IMiD), bortezomib (PI) and Dex, improved depth of response and progression-free survival (Moreau 2019). Dex is also used in all treatment options for patients with relapsed and /or refractory MM in combination with the second-generation IMiDs lenalidomide and pomalidomide and PIs carfilzomib and ixazomib (Gariani 2018). Despite spectacular therapeutic improvement, few patients are cured, therefore it is necessary to better understand the precise mechanisms of action of each agent alone or in combination.

Dex exerts its biological functions by binding to the glucocorticoid receptor (GR) encoded by *NR3C1*. Upon Dex binding, the complex translocates to the nucleus, where it associates with DNA at GR binding sites, acts as a transcription factor (TF) and regulates gene expression (Reddy 2009). GR binding appears to be pre-programmed by the binding of lineage-specific TFs and chromatin accessibility prior to exposure (Biddie 2011; John 2011). At these loci, GR cobinds with cell-specific pioneer TFs including C/EBP in the liver (Grontved 2013), PU.1 in the macrophage lineage (Oh 2017), and AP1 in murine hepatocytes (Biddie 2011). GR binds predominantly at distal enhancers (Reddy 2009) and drives transcription by interacting with gene promoters via chromosomal loops. GR binds thousands of locations across the genome but only few enhancers cooperate with each other to activate Dex-responsive genes (Vockley 2016; Mcdowell 2018).

However, how these enhancers combine to induce gene expression is still poorly understood. A previous study exploiting protein-directed chromatin interactions approach suggests that at GR-responsive genes, chromatin interaction loops between enhancers and promoters are pre-established while in a subset of genomic loci, GR binding induces de novo interactions (Kuznetsova 2015). A recent study showing high-resolution genome-wide maps of chromatin interactions in response to Dex confirms and extends the model that GR binding acts predominantly through pre-determined chromatin interactions and increases their frequency (D’Ippolito 2018). However, these studies did not resolve the influence of increased chromatin accessibility on chromatin loops and enhancer activity as shown by Staverva et al (2015) (Stavreva 2015), as well as its consequences at the functional level.

Given that Dex is an essential drug in the treatment landscape of MM disease course, the analysis of its molecular action on the genome of myeloma cells and the consequences on transcriptional heterogeneity are needed in order to better understand treatment escape.

## Results

### GR binds to a pre-programmed landscape in malignant plasma cell

To identify the genomic features associated with GR binding in malignant plasma cells, we firstly defined the pre-existing chromatin landscape at GR binding sites. To this end, we performed chromatin immunoprecipitation sequencing (ChIP-seq) for GR, ChIP-seq histone marks for H3K27ac, assay for transposable-accessible chromatin sequencing (ATAC-seq) and RNA sequencing (RNA-seq) in the Dex-sensitive human myeloma cell line MM.1S exposed to Dex (0.1µM) or EtOH for 1 hour (Fig. 1a). We found that 46% (8,476/18,444) of the GR binding sites were located in promoter distal regions based on their 5kb-distance to the transcription start site (TSS) (Fig. 1b). We next defined the chromatin states for GR binding sites prior to treatment by performing chromatin hidden Markov modeling (chromHMM) (Fig.1b, Additional file 1: Fig. S1a). As expected, this analysis revealed a large majority (86%) of GR binding sites at active and accessible regulatory regions including strong enhancers and strong promoters marked with H3K4me1, H3K4me3, H3K27ac and open chromatin as exemplified at *GILZ* (*TSS22D3*) locus (Additional file 1: Fig. S1b) and a small subset (3%) of GR binding sites at weak enhancers marked with histone mark H3K4me1, not marked with H3K27ac and mostly DNAse I inaccessible. Chromatin-state assignments were confirmed by our H3K27ac ChIP-seq and ATAC-seq analyses in MM.1S (Fig.1c). Stretch-or super-enhancers (SEs) are clusters of enhancers spanning multiple kb of DNA which exhibit high density of H3K27ac (Hnisz 2013; Whyte 2013). We observed GR binding sites in 99% (808/815) of SEs identified in MM.1S (Loven 2013) (Additional file 1: Fig. S1c). Overall, these results confirmed that GR binds multiple active enhancers individually or cooperating as SE.

**Figure 1.**
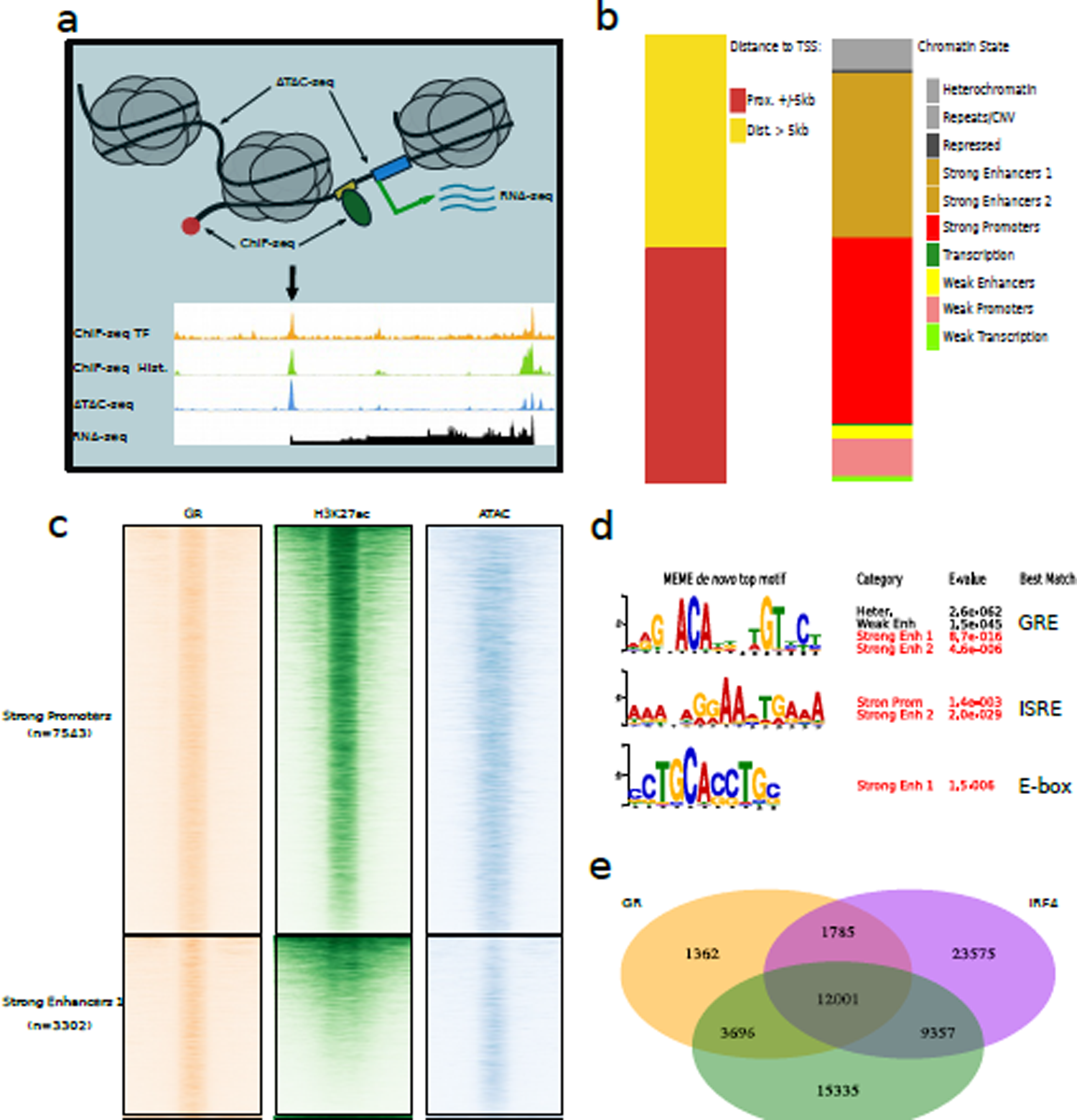
Pre-programmed chromatin landscape guides GR binding in malignant plasma cells. **(a)** Scheme of sequencing data used to define chromatin landscape of MM.1S. **(b)** Genomic annotation of the 18,844 Dex-specific GR binding sites according to TSS from reference genome hg19 (left) and according to MM.1S functional chromatin states (right), see chromHMM annotations in Figure S1a. **(c)** Heatmaps of GR binding, H3K27ac marks and ATAC-seq signal in a 4kb-region centered on GR binding site. GR binding sites are divided according to chromatin states defined as pre-active regulatory regions (Strong promoters, Strong enhancers 1 and Strong enhancers 2) and pre-inactive regulatory regions (Weak promoters, Weak enhancers and Heterochromatin) and ordered on the y-axis based on the mean normalized GR value. **(d)** De novo MEME top motif enriched in MM.1S chromHMM functional states. **(e)** Overlap of ChIP-seq peaks for GR (MM.1S Dex 1 hour), IRF4 (MM.1S EtOH, loven 2013) and H3K27ac (MM.1S EtOH). **(f)** Radar chart showing GR binding partners using rapid immunoprecipitation mass spectrometry of endogenous protein (RIME) method in MM.1S (Dex 1 hour); Arbitrary Units.

We then investigated pre-existing TFs occupancy at GR binding sites. A de novo enrichment analysis showed glucocorticoid responsive elements (GREs) in strong enhancers (1 and 2) and revealed IRF4 motifs in strong enhancers 2 chromatin state (Fig.1d). IRF4 is the master regulator of aberrant regulatory network in myeloma cells (Shaffer 2008). Furthermore, we observed a strong overlap of GR binding sites, IRF4 occupancy and H3K27ac peaks (Fig. 1e). Interestingly, among the IRF4 binding sites shared by Dex-activated GR binding sites, only a subset of sites were enriched for GRE (18%; 2,502/13,786) or half GRE (7%; 1,031/13,786). Enrichment analysis of those sites indicated that GR binding sites with GRE or half GRE were associated with biological processes related to Dex function in myeloma cells, notably apoptotic signaling pathway (Additional file 1: Fig. S2). These results are in line with previous studies showing the importance of GRE in Dex response (Vockley 2016).

Next, a rapid immunoprecipitation mass spectrometry of endogenous protein method was conducted to identify GR binding partners in MM.1S (RIME; Mohammed 2016; Additional file 2: Table S1; Fig. 1f). As anticipated, IRF4 was one of the top ranking partners along with TFs including IKZF1 and IKZF3 that are essential in multiple myeloma (Kronke 2014) and MEF2C that is important for *MYC* SE activity and expression in primary effusion lymphoma (Wang 2020). Unlike IRF4, we did not find cognate DNA-binding motifs close to the GR binding sites for IKZF1/3 and MEF2C; this raises the possibility that these TFs might exert roles in trans on GR transcription programs analogous to the effects of the MegaTrans complex on the ER α-regulated functional enhancers (Liu 2014). GR interacts also with the catalytic ATPase BRG1 (SMARCA4) and several associated subunits, including BAF170 (SMARCC2), BAF155 (SMARCC1), BAF60B (SMARCD2) of the SWI/SNF chromatin remodeling complex capable of moving and displacing nucleosome.

Together, those results confirm the importance of pre-programmed chromatin landscape in guiding most of GR binding at open and active genomic loci and lowly GR-bound at latent regions. Furthermore, IRF4 contributes to predetermine GR binding. Finally, GR could act via a complex similar to MegaTrans complex of ER, composed of several lineage-specific TFs, the remodeling complex SWI/SNF and potential coactivators to regulate Dex-responsive genes. This complex could be assembled in trans on GR-bound functional enhancers and could require the presence of GR (Additional file 1 - Fig.S3).

Since there are many more regulatory regions that bind GR than Dex-responsive genes, it is important to draw a functional map of GR-activated enhancers to better understand Dex action in myeloma cells.

### Landscape of enhancer cluster cooperation in malignant plasma cells responsive to Dex

Given that enhancer activity modifications are associated with gene expression changes (Mcdowell 2018), we firstly characterized the GR-bound regions that showed significant increase of H3K27ac upon Dex exposure to identify functional enhancers of target genes in myeloma cells (Fig.2a; Additional file 3: Table S2). This subset of regions which contained a small percentage of GR binding sites (5.3%, 864/16,228; see Materials and Methods) occurred predominantly (85%) in distal enhancers located more than 5kb away from closest TSS (Fig. 2b).

**Figure 2.**
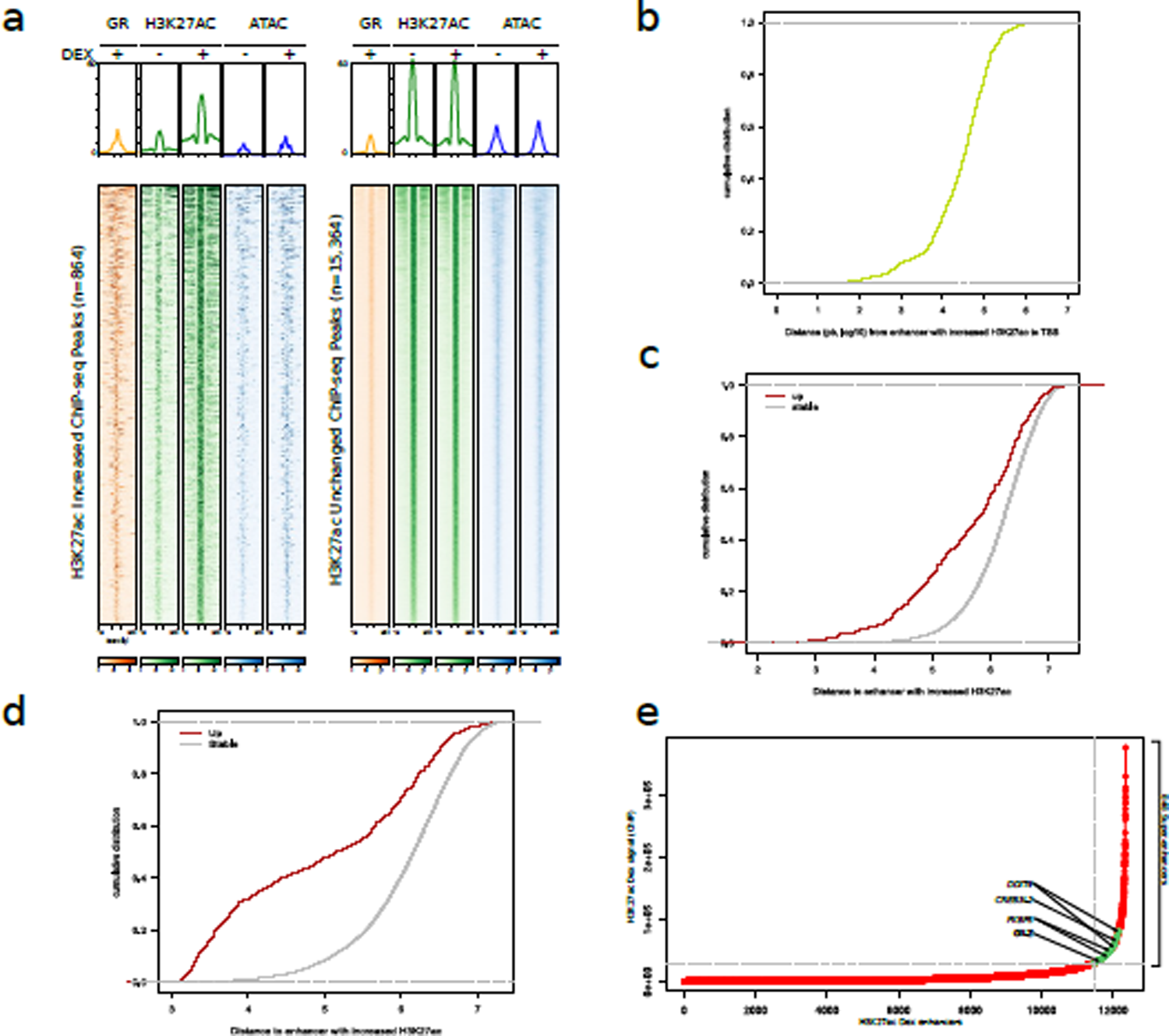
Characterization of functionnal enhancers in MM.1S upon Dex exposure. **(a)** Heatmaps of GR binding in MM.1S exposed to Dex (1 hour), H3K27ac marks MM.1S (EtOH and Dex, 1 hour) and ATAC-seq signal MM.1S (EtOH and Dex, 1 hour) in a 4kb-region centered on GR binding site. Left panel are regions with increased H3K27ac signal after Dex exposure compared to EtOH, right panel are regions with stable H3K27ac signal. **(b)** Cumulative distribution plot of relative distance from H3K27ac Dex-increased enhancers to TSS (log10(bp)). **(c)** Cumulative distribution plot of relative distance from H3K27ac Dex-increased enhancers for Dex up-regulated genes (red) and Dex stable genes (grey) (log10(bp)). **(d)** Cumulative distribution plot of relative distance between H3K27ac Dex-increased enhancers (red) and between H3K27ac Dex-increased enhancers and H3K27ac Dex-stable enhancers (grey) (log10(bp)). **(e)** SEs determination using H3K27ac ChIP-seq peak signals of MM.1S exposed to Dex 1 hour. Associated Dex-responsive genes are indicated by an arrow. Dex-specific Super-Enhancers are colored in green.

Interestingly, the up-regulated enrichment of H3K27ac at GR-bound regions was associated with increased chromatin accessibility compared to GR-bound regions without H3K27ac occupancy changes (Fig. 2a). The functional significance of the active enhancers that gained H3K27ac was evidenced by the fact that upregulated genes were nearer to them than other genes (Fig.2c) and that they were closer to each other than other active enhancers (Fig. 2d). Importantly, the ROSE algorithm (Whyte 2013) distinguished 121 Dex-specific SEs. Nearest gene analysis identified known GR-responsive genes, including *DDIT4*, *CREB3L2*, *FKBP5* and *GILZ* (Fig.2e). Thus indicating that individual enhancers that gained H3K27ac after Dex exposure may cluster together and reach the SE status. In addition, their linear proximity could promote their cooperativity for a strong and rapid expression of their target genes. However, individual enhancers may also spatially cooperate and loop to genes further away (Petrovic 2019; Mumbach 2017). Therefore, to evaluate the regulatory elements looping patterns, we generated high-resolution contact maps of active enhancers and target genes by H3K27ac HiChIP in MM.1S cells (Fig. 3a). We identified 21,249 and 23,278 H3K27ac chromatin interactions across the genome in MM.1S EtOH and MM.1S Dex, respectively. As anticipated, H3K27ac ChIP-seq signal was enriched at anchor regions (Additional file 1: Fig. S4a). Interestingly, the median number of interactions was higher in Dex-treated MM.1S (3 vs 2) however the median interaction distance was similar (124 kb for MM.1S EtOH vs 127 kb for MM.1S Dex) (Additional fille 1: Fig. S4b). The majority of H3K27ac-associated interactions occurred between proximal-proximal (proximal<5kb from the closest TSS) interactions or proximal-distal (distal>5kb from the closest TSS) interactions (Additional file 1: Fig. S4c). GR binding was preferentially found at looping anchors (Additional file 1: Fig. S4d).

**Figure 3.**
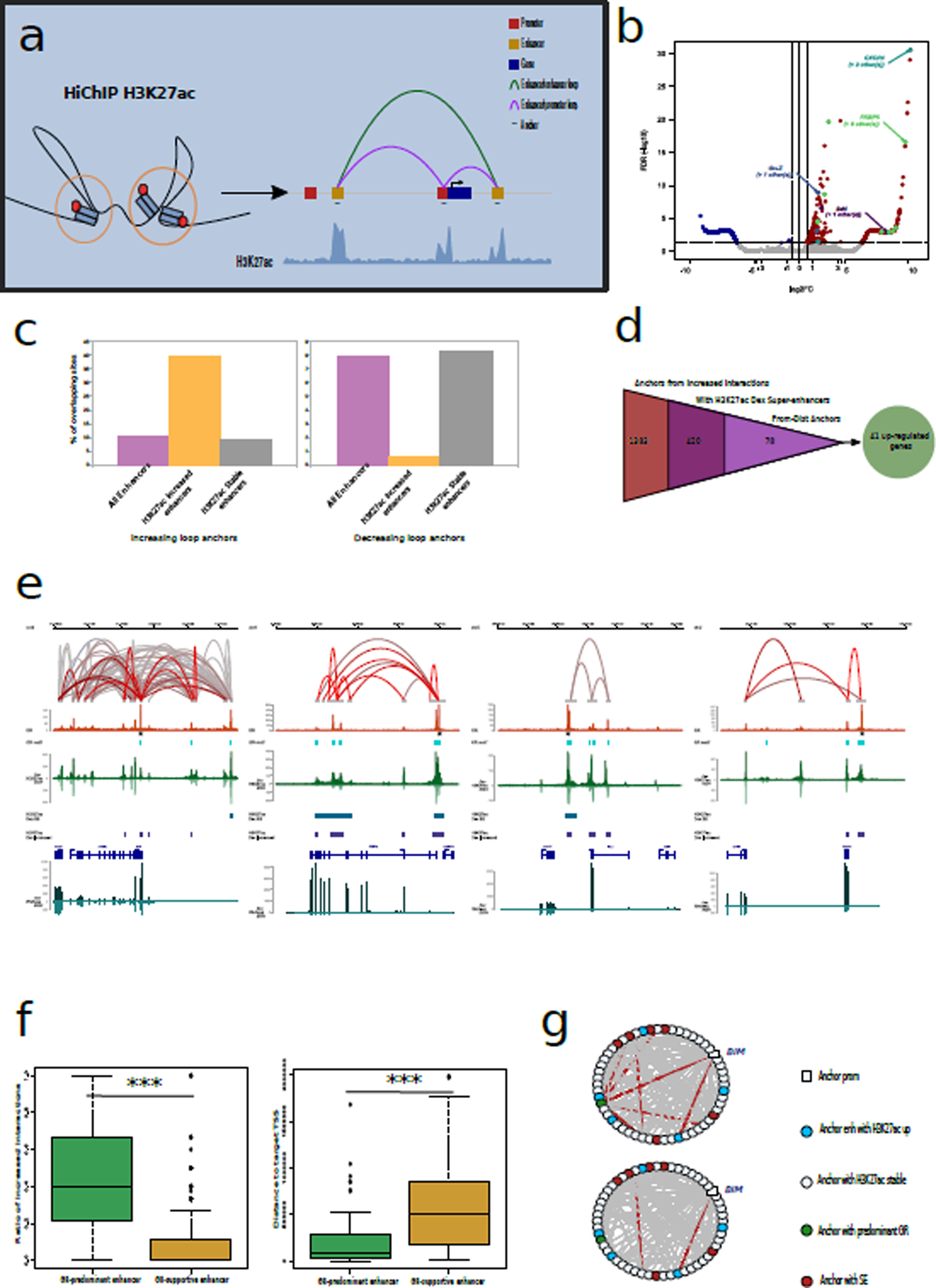
GR binding induces enhancer cooperativity. **(a)** Scheme depicting H3K27ac loop interactions between regulatory regions through linear genome. **(b)** Volcano plot of HiChIP H3K27ac differentially induced (FDR<0.05) loops between EtOH and Dex MM.1S cells, increased (log2FC>0.6) loops in red and decreased (log2FC<-0.6) loops in blue. **(c)** Overlapping of enhancers with anchors linked to Dex-increased loops (left) or Dex-decreased loops (right). **(d)** Diagram illustrating the overlap of Dex super-enhancers with Dex-increased loop anchors and anchors controlling up-regulated genes. **(e)** Snapshots for *BIM*, *FKBP5*, *GILZ* and *CXCR4* loci illustrating example of GR binding to its consensus motif GRE that increases the H3K27ac ChIP-seq signal as much as the pre-existent H3K27ac chromatin interactions, predominant enhancer is indicated by a black star; interactions loops are colored according to their respective log Fold Change (LogFC) from deep blue (LogFC=-10) to deep red (LogFC=10), LogFC around zero being greyed out. **(f)** Box plot illustrating the ratio of Dex-increased chromatin loops for the anchor containing a GR-predominant enhancer (green) or the anchors containing GR-supportive enhancers (yellow; left) and the distance from closest Dex-induced TSS of GR-predominant enhancer and GR-supportive enhancers (right). *** p<0.001. **(g)** Circos plots depicting interactions network of *BIM* gene, (top) complete regulatory network, (bottom) network without the interactions linked to predominant enhancer (green).

Comparison of H3K27ac interaction loops between MM.1S Dex and MM.1S EtOH revealed that GR binding significantly increased chromatin interaction for 917 loops (4.3%) and significantly decreased interaction for 581 loops (2.7%). GR binding was significantly associated with Dex-increased H3K27ac HiChIP loops (Additional file 1: Fig. S5). These loops were enriched in distal-distal interactions and depleted in proximal-proximal interactions compared to stable or decreased loops (Additional file 1: Fig. S6). The CTCF has been shown to facilitate and stabilize distal-proximal interactions loops (Ren 2017; Kubo 2021). In line with previous studies, our results of CTCF ChIP-seq showed that CTCF binding was unchanged in Dex-treated cells compared to EtOH-treated cells and that CTCF occupancy was lower in increased loop anchors compared to stable interactions suggesting that chromatin interactions that are more dynamic in response to Dex may be bound by other TFs (Additional file 1: Fig. S7) (D’ippolito 2018).

Next, we focused on loops that significantly increased their frequency upon Dex treatment. Analysis of distal-proximal loops identified the essential gene for Dex-induced death in MM.1S *BIM* (*BCL2L11*), the chemokine receptor gene *CXCR4* known to be associated with MM progression and poor prognosis and ubiquitously Dex-responsive genes including *GILZ* and *FKBP5* (Fig.3b). Notably, Dex-increased loops displayed a strong sequence overlapping (∼40%) with H3K27ac-increased enhancers and low overlapping (8%) with the stable enhancers while this was the reverse, albeit to a lesser extent, for Dex-decreased loops (Fig. 3c). As described above, individual active enhancers collaborate to form linear clusters or SEs. We showed that 420 SEs overlapped with increased loop anchors. Among those, distal-proximal anchors were linked to 41 Dex-responsive genes including *BIM*, *GILZ* and *FKBP5* (Fig. 3d, Additional file 4: Table S3). We also showed that Dex-increased interaction loops involved individual enhancers as well as SEs that coalesced to form a spatial network of interactions with a single target promoter, termed enhancer cliques by Mumbach et al (Mumbach 2017) (Fig. 3e). Notably, we observed two types of responsive genes, those within highly connected cliques such as *BIM* or *FKBP5* and those that spatially interacted with a lower connectivity such as *GILZ* or *CXCR4*.

We next investigated how these enhancers cooperate. We observed that among the enhancers of the same gene interaction network there was always one which had a higher GR peak than the others and that this peak was most often associated with a significant enrichment of GRE (indicated by a star in the representative examples of Fig.3e), thus suggesting a functional hierarchy among enhancers in which a GR-predominant enhancer (predominant enhancer) collaborates with GR-supportive enhancers (supportive enhancers) as evidenced for estrogen α binding sites (Carleton 2017). To test this, we firstly determined the predominant enhancer for each interaction network (loop anchors) of the Dex-responsive genes (Additional file 5: Table S4; see Materials and Methods). Predominant enhancers were characterized by a higher degree of interactions and a greater proximity to the Dex-responsive gene TSS compared to the supportive enhancers (Fig. 3f). In addition, not tracing interactions connected to the predominant enhancer dramatically lowers the clique connectivity (Fig. 3g, Additional file 1: Fig. S8).

In summary, the results suggest that GR binding induces enhancer cooperativity to form enhancer cliques as well as linear clusters to drive transcription. Moreover, it appears that a functional enhancer hierarchy with one predominant enhancer may exist in these high-order structures.

### Dex increases chromatin co-accessibility of distal and proximal regulatory regions

Since we identified a subset of functionnal enhancers that gained both H3K27ac marks and chromatin accessibility upon Dex exposure (Fig.2a), we investigated the role of GR binding in chromatin remodeling and its potential impact on spatial enhancer activity. ATAC-seq analysis revealed a strong correlation between increased H3K27ac chromatin loops and gain of chromatin accessibility (p<0.0001; Fig. 4a, Additional file 6: Table S5). Notably, these regulatory regions with increased chromatin accessibility overlapped both SEs and individual enhancers (Fig.4b) and showed a higher GR binding than regions with stable ATAC abundance (Fig.4c). Furthermore, de novo motif analysis revealed that GRE was enriched in Dex-increased ATAC-seq peaks (Fig.4d) while IRF4 binding motif (ISRE) was found in stable ATAC (Fig.4e). Altogether, these results demonstrate that Dex treatment leads to chromatin accessibility gain at the increased H3K27ac loop anchors and raise the possibility of a higher level Dex-responsive genes regulation depending on co-accessibility of regulatory regions involved in enhancer cliques.

**Figure 4.**
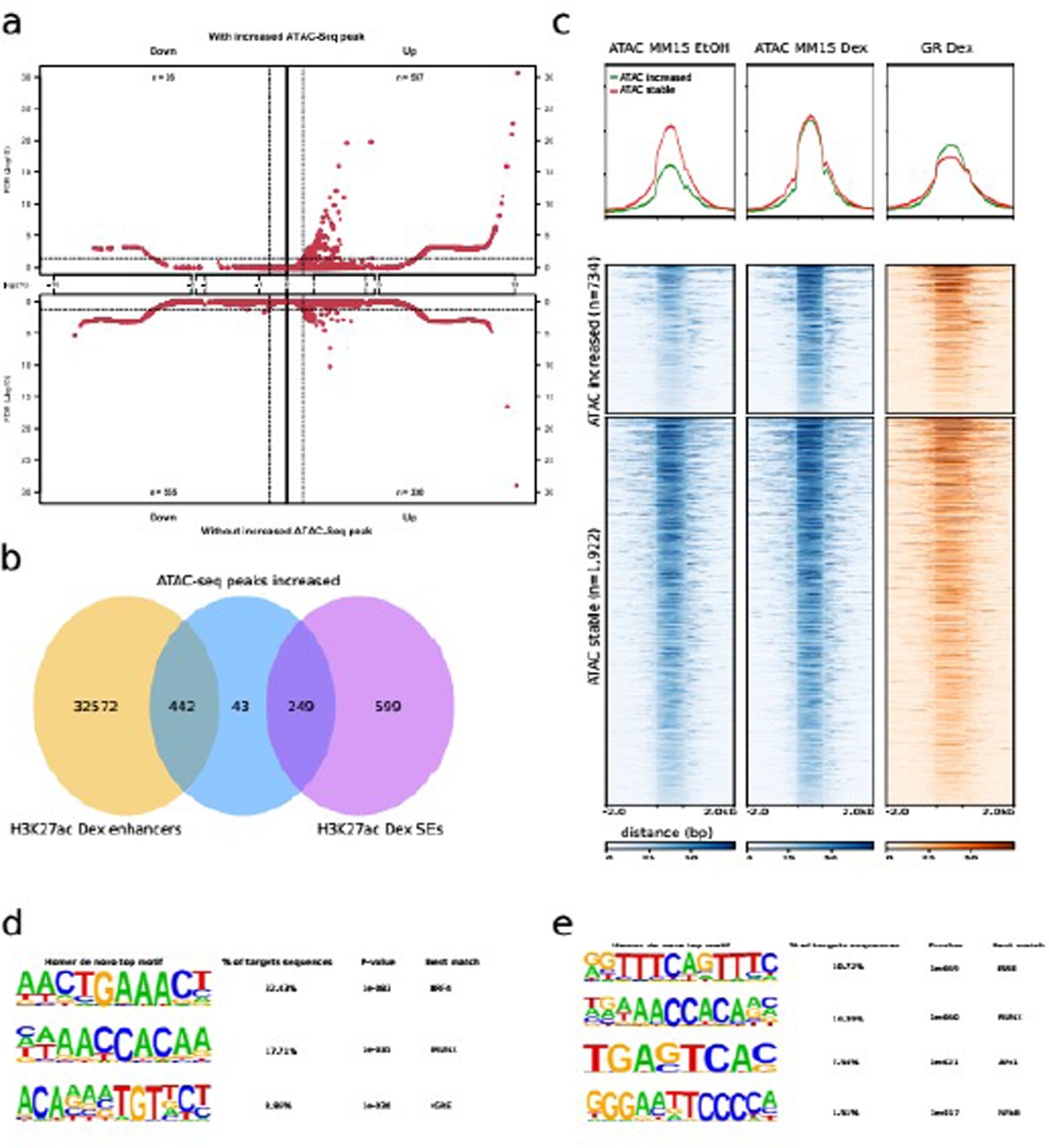
Effect of GR binding on chromatin accessibility in anchors of Dex-increased H3K27ac HiChIP loops. **(a)** Volcano plot showing the differential H3K27ac chromatin interactions within Dex-increased ATAC-seq peaks (top) or without (bottom, mirrored). **(b)** Venn diagram of overlap of Dex-increased ATAC-seq peaks located within anchors of increased H3K27ac interactions with SEs and individual enhancers. **(c)** Heatmaps illustrating ATAC-seq signal (EtOH and Dex conditions) and GR ChIP-seq signal in ATAC-seq Dex-increased or Dex-stable peaks located within anchors of Dex-increased H3K27ac HiChIP loop. **(d-e)** *De novo* motif discovery on Dex-increased ATAC-seq peaks **(d)** or Dex-stable ATAc-seq peaks **(e)**.

To test this, we performed simultaneous profiling of open chromatin and gene expression from the same cell (scMultiome) across 5,000 nuclei extracted from MM.1S exposed to Dex or EtOH for 1 hour and 4 hours. After normalization, all data were subjected to Uniform Manifold Approximation and Projection for Dimensional Reduction (UMAP) (Becht 2019). Cell clusters were computed and color-coded according to treatment. We found that Dex altered chromatin accessibility in agreement with bulk ATAC analysis and that prolonged Dex exposure to 4 hours did not modify accessibility landscape (Fig.5a left). We then focused on the GR predominant and supportive enhancers (Additional file 4: Table S3). We observed a significant increase of chromatin accessibility in these regulatory regions, however the gain was more pronounced in the predominant enhancers compared to supportive enhancers (Fig. 5a middle, right). In the same way, after Dex exposure, the number of cells with accessible predominant or supportive enhancers was significantly increased although the mean difference was more pronounced for the predominant enhancers (Fig.5b). Then, we applied Cicero (v1.3.4.11) (Pliner 2018) to calculate the co-accessibility scores for each predominant enhancer. As anticipated, co-accessibility changes between predominant enhancers and supportive enhancers or promoters were higher after Dex exposure (p=0.02, Fig. 5c). Next, we focused on gene-specific networks that gained co-accessibility (21/62) after Dex exposure, among them we found the universally Dex-induced genes *FKBP5* and *GILZ* and myeloma-specific inducible genes *BIM* and *CXCR4*. In the *FKBP5* gene locus, GR binding led to a significant increase of co-accessibility within individual elements of each SE and between SEs (Fig.5d, dashed boxes) in agreement with bulk H3K27ac HiChIP signals (Fig.3e). Notably, the predominant enhancer opens significatively as well as 3 other regions (Fig.5e). The network graph of the predominant enhancer, supportive enhancers and promoter of *FKBP5* showed the increased complexity of interactions between the predominent enhancer and the other regulatory regions as well as the central position of the predominant enhancer after Dex exposure (Fig.5f). As for *FKBP5* gene locus, GR binding at the *GILZ* locus led to a significant increase of co-accessibility within individual elements of the SE, and also between predominant enhancer, supportive enhancer and the promoter (Additional file 1: Fig.S9 a and b). Notably, the predominant enhancer opens significantly as well as another region (Additional file 1: Fig.S9c). Regarding *BIM* and *CXCR4*, Dex exposure provoked the co-accessibility increase between the predominant enhancer and the promoter (Additional file 1: Fig.S10 a and b; S11 a and b). However, at *BIM* locus, only the predominant enhancer opens significantly while at *CXCR4* locus, both predominant enhancer and promoter open significantly (Additional file 1: fig S10c, S11c).

**Figure 5.**
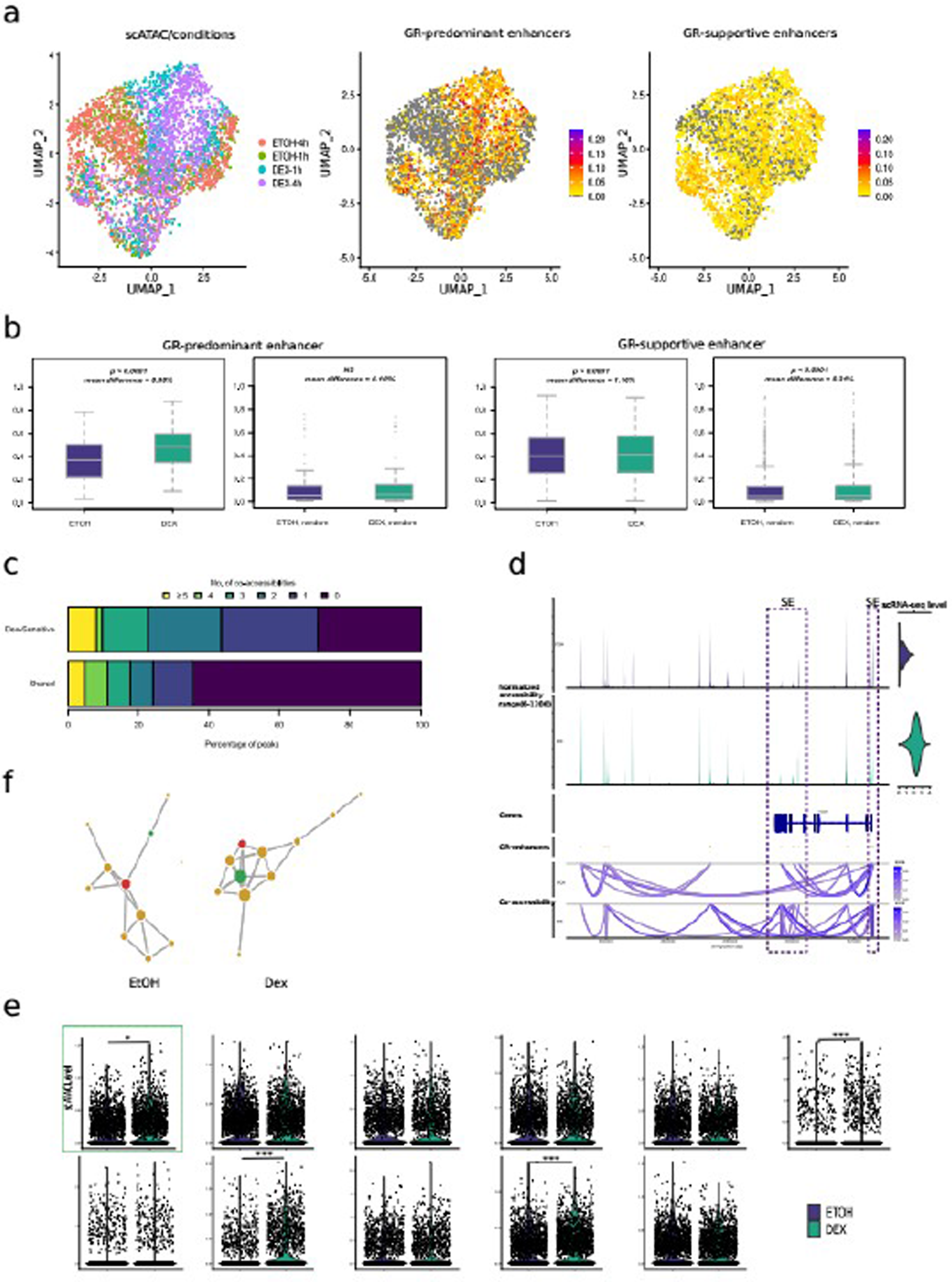
Dex increases chromatin co-accessibility of distal and proximal regulatory regions. **(a)** Uniform Manifold Approximation and Projection (UMAP) of scATAC-seq profiles in MM.1S cells, colored by sample of origin (left), GR-predominant enhancers activity score (middle) and GR-supportive enhancers activity scores (right). **(b)** Boxplots illustrating percentages of cells with open GR-predominant enhancer vs. the same number of randomly selected regions (left) and open GR-supportive enhancer vs. the same number of randomly selected regions (right). **(c)** Barplot showing distribution of number of co-accessibilities shared between the 2 conditions or Dex-Sensitive. **(d)** Snapshot of *FKBP5* locus revealing aggregated chromatin accessibility for EtOH (blue) and Dex (green) conditions, refseq annotation of *FKBP5*, localization of the ATAC peaks overlapping with GR-regulatory regions (green: GR-predominant enhancer; red: *FKBP5* promoter; yellow: GR-supportive enhancers), co-accessibility links between the selected peaks in EtOH and Dex conditions. *FKBP5* scRNA-seq expression level violin plots (top-right). **(e)** Violin plots illustrating accessibility of each peak across the treatment conditions, statistically significant differential accessibility is shown (* adjusted pval<0.05, *** adjusted pval<0.001). **(f)** Network graph representing GR-regulatory regions of *FKBP5*, edges width reflects co-accessibility score, node size depends on the number of connections to other nodes. Same colors as in Fig. 5d were used.

Together, these data showed that the GR binding increases the chances of having a co-accessibility between the predominant enhancers and the other regulatory regions.

### Cell-to-cell transcriptional heterogeneity within myeloma cells after Dex treatment

Since we showed that the predominant enhancer cooperates with supportive enhancers and the promoter of the Dex-responsive genes and that Dex exposure increases co-accessibility of these regulatory elements, we next evaluated the consequences of Dex-induced gene regulatory network cooperativity on transcriptional heterogeneity. To do this, we performed scRNA-seq analysis in MM.1S cells collected at 4 and 24 hours in presence of Dex (0.1µM) or EtOH. We focused our analysis on the genes most strongly induced by Dex. Analysis of logFC distribution permitted to isolate 51 highly induced genes, termed single cell Dex-activated genes (scDAGs) (Additional file 1; Fig. S12a). All scDAGs except 2 genes were already in the three datasets (GR ChIP-seq, H3K27ac HiChIP and bulk RNAseq, Additional file 4: Table S3). In order to measure the level of cell-to-cell transcriptional variability, we calculated scDAGs expression information entropy (Pastore 2019; Landau 2014) and found that among cells exposed to Dex, the median expression increased, leading to an increased fraction of positive cells (fpc), toward 1; hence we observed a decrease in entropy, i.e. less heterogeneity among the cells. However, gene expression among the cells remained heterogeneous, with a high inter-quartile range (IQR) (Fig. 6a). To further examine the transcriptional variability that remained after Dex exposure, we analyzed the relationships between scDAGs. Although not very strong, we observed a higher two-by-two correlations between scDAGs than between random genes (Additional file 1: Fig. S12b-d). Interestingly, scDAGs were clustered in two main groups, a large cluster (cluster 1; 45/51 genes) including the pro-apoptotic gene *BIM* and ubiquitous GR-responsive genes such as *GILZ*, *FKBP5* and *DDIT4* and a second cluster encompassing 6 genes including *CXCR4* (Fig. 6b). These results suggest that within Dex-treated cells, two subpopulations coexist. A cell population that predominantly expresses a large majority of scDAGs (referred as highly Dex-responsive cells) and another population of cells expressing a reduced number of scDAGs.

**Figure 6.**
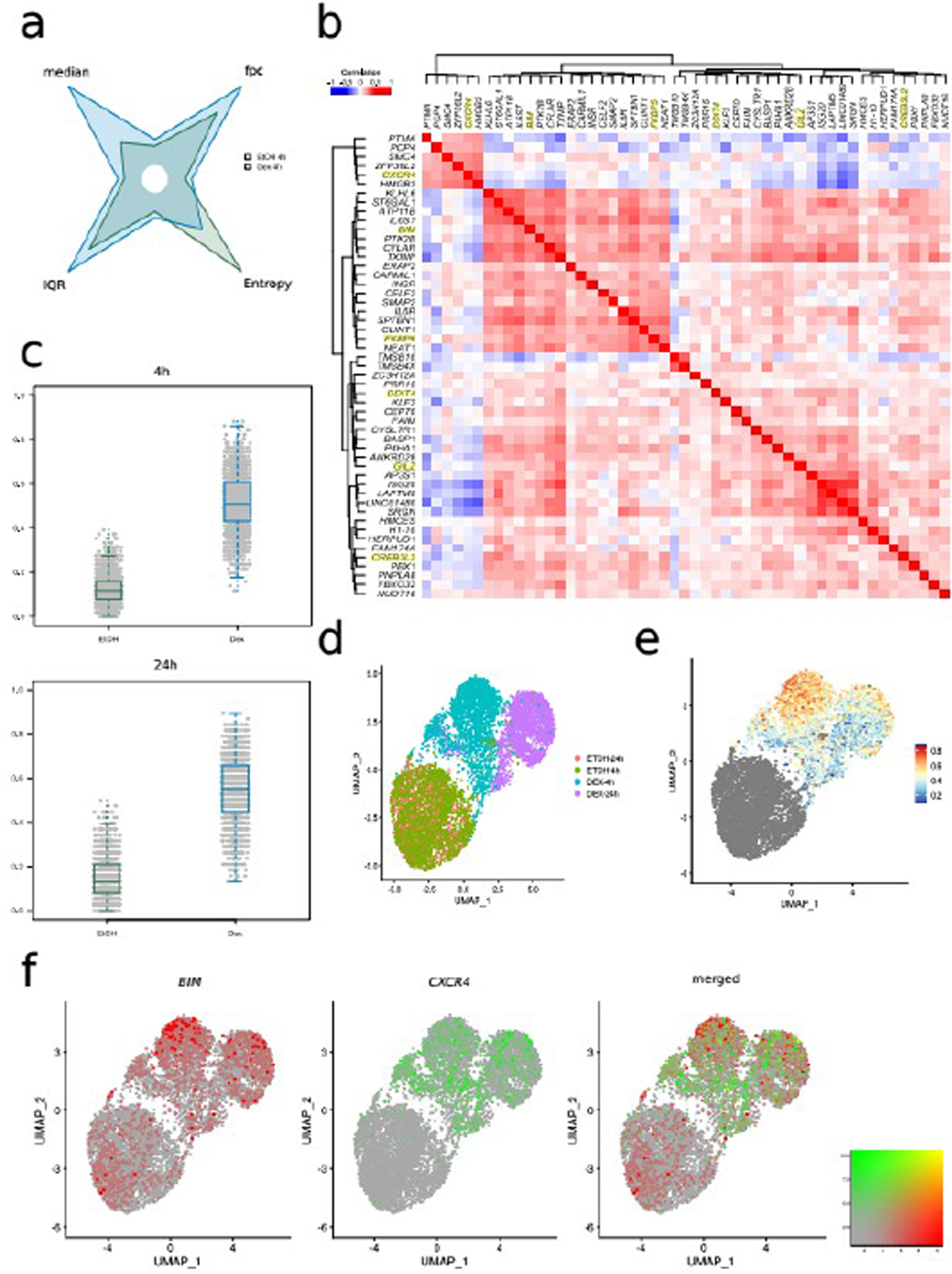
Cell-to-cell transcriptional heterogeneity after Dex exposure. **(a)** Shuriken plot illustrating median, IQR, fpc and entropy for control and Dex-treated cells; for each parameter, maximum value is used as reference. **(b)** Heatmap of two-by-two correlations among the 51 DAGs. **(c)** Ratio of Responding Genes after 4 hours (top) and 24 hours (bottom) of treatment. **(d)** UMAP plot of MM.1S cells from scRNA-seq colored by condition. **(e)** UMAP plot colored by the ratio of responding genes for 4 hours of exposure. **(f)** UMAP plots colored for *BIM* expression (red), *CXCR4* (green) expression and merged.

To test this, we firstly employed the method recently described by Hoffmann et al. (Hoffman 2020), to estimate the number of scDAGs expressed in each cell after 4 and 24 hours of Dex or EtOH exposure (Fig. 6c, Additional file 1: Fig. S13). We found that an EtOH-treated cell had a median background ratio of responsive genes (RRG) of 12% (6 /51 scDAGs) and a Dex-treated cell had a median RRG of 51% (26/51 scDAGs) similar to that of 24h Dex exposure (55%; 21/38 DAGs). Then, we visualized the scDAGs transcriptional variability by performing dimensional reduction using UMAP. As anticipated, UMAP reduction separated perfectly cells according to treatment conditions (Fig. 6d). Finally, we colored map the RRG on the UMAP plot (Fig. 6e). Interestingly, although there was an important cell-to-cell heterogeneity, highly Dex-responsive cells tended to cluster together while poorly responsive cells were scattered around. Merged UMAP plots colored according to gene expression of uncorrelated genes *BIM* and *CXCR4* (R2 = −0.02) clearly showed that in most of cells expression of *BIM* and *CXCR4* was mutually exclusive (Fig. 6f). The same applies to *CXCR4* and *GILZ* and *CXCR4* and *FKBP5* (Additional file 1: Fig. S14).

Altogether, scRNA-seq analysis revealed that on the average MM.1S cells expressed only half of the overall Dex-responsive genes. In addition, in the vast majority of poorly responsive cells (i.e. RRG<50%), *BIM*, the most important GC-induced death gene in MM, was not expressed.

## Discussion

Although the efficacy of Dex in MM can largely be attributed to GR-induced apoptosis, the genomic responses to Dex treatment in malignant plasma cell genome remain unknown. Given that Dex is used at all stages of treatment, it was crucial to investigate its molecular mode of action by using new genomic tools in order to better understand treatment escape and provide new insights into combination therapy options.

In this study, we confirmed that the plasma cell-specific epigenomic landscape was primed by the lineage-specific pioneer factor IRF4 and that GR binds pervasively to a largely preprogrammed landscape of active enhancers. However, despite a strong association of H3K27ac with GR binding within enhancers engaged in long-range interactions to form lineage-specific networks, the changes in enhancer activity upon Dex exposure are limited to a small number of distal elements. These elements contribute to the regulation of GR-responsive genes in a combinatorial manner. Indeed, our study showed that GR binding modifies H3K27ac across large stretches of enhancers termed SEs. These dynamic SEs are associated with Dex target gene promoters by chromatin interactions and ensure strong gene expression. Moreover, our high-resolution regulatory connectivity maps revealed that GR binding promotes also an increase in the frequency of regulatory DNA loops linking multiple distal enhancers to their target genes previously reported as enhancers cliques (Mumbach 2017). This mode of action is similar to that already described for other TFs (Wang 2014; Petrovic 2019). Overall, our results demonstrate that specific combinations of enhancers collaborate to produce a proper and selective Dex response. Among these enhancers, we showed the importance of GRE motif and distance to TSS for GR binding in agreement with previous studies (Vockley 2016; Mcdowell 2018). Together, these data suggest the existence of a hierarchy within enhancer clusters with a predominant site and supportive sites which collaborate following GR binding as it has been previously reported in other lineages (Shin 2016; Carleton 2017; Saravanan 2020; Huang 2021). Recent work in acute lymphoblastic leukemia has shown that a specific enhancer is necessary to mediate interaction with the promoter resulting in *BIM* activation (Jing 2018). In the present study, we also showed that this enhancer is involved in most of dynamic up-interactions in Dex response. We can assume that this enhancer is the regulatory element which collaborates in a super-additive fashion with other enhancers to achieve full expression of *BIM*. Targeted genome engineering could be used to directly test this hypothesis and determine the functional contribution of the other enhancers (Thomas 2021). We can speculate that these enhancers may work together to increase the local protein concentration and form phase-separated transcriptional condensate to rapidly activate gene expression as previously observed for other nuclear receptors (Boija 2018; Nair 2019; Saravanan 2020).

As we found that chromatin accessibility correlates with the frequency of enhancer interactions with each other or nearby gene targets, we integrated scATAC-seq and scRNA-seq with HiChIP and ChIP-seq datasets in order to study the influence of chromatin accessibility to the enhancer activity. Our results showed that Dex treatment leads to an increase of the number of open regions, in particular for predominant enhancers leading to a significant increase in the co-accessibility between this enhancer and other regulatory regions. Altogether, these results suggest that after Dex treatment a cell is more likely to have the optimum connections for the correct expression of a target gene and that the predominant enhancer acts as a regulatory hub between promoters and other supportive enhancers as previously described for androgen receptor (Huang 2021).

Even if additional studies are needed to analyze chromatin heterogeneity among individual cells by combining single cell conformational studies (ChIA-Drop; Zheng 2019) with single cell epigenomics (CUT&Run and CUT&Tag protocols; Hainer 2019; Kaya 2019), our study demonstrates the importance of taking into account all enhancers involved in spatial gene regulation in order to better understand the mode of action of TFs.

Finally, we showed that variability in chromatin accessibility was associated with a heterogeneous response to Dex in myeloma cells as it has been reported before for breast cancer (Hoffman 2020). Indeed, we found a cell-to-cell variability regarding the expression of Dex-responsive genes, firstly on the average each cell expressed approximately half of the Dex-induced transcriptional network and secondly a subset of cells did not express *BIM*, a major mediator of Dex-induced apoptosis in myeloma cell lines (Kervoelen 2015). Our results highlight the complex interplay between cell-to-cell modulation of chromatin accessibility and distal-proximal and distal-distal interaction looping increase upon GR binding that could lead to a mutually exclusive expression of *BIM* and *CXCR4* and provide new insights into the mechanisms of drug escape while considering that GR levels can be a limiting event in Dex treatment (Kervoelen 2015; Heuck 2012) (Additional file 1: Fig. S15).

Interestingly, our results showed that IKZF1 and IKZF3 are among the few GR-cobond partners, suggesting that these lineage specific TFs could play a role in Dex response as previously described for the MegaTrans complex in the functionally active estrogen-regulated enhancers (Liu 2014). A recent study showed that these TFs are degraded by IMIds (Sievers 2018). Since both drugs are combined to treat MM patients, we cannot exclude an antagonistic role of these molecules, further studies to identify the GR-IKZF1/3 target genes if any are warranted.

Given the potential role of *CXCR4* in tumor growth and dissemination (Alsayed 2007; Roccaro 2015), its increased expression upon Dex exposure in a subset of MM.1S cells that do not express the proapoptotic gene *BIM* raises the provocative possibility that minor populations of myeloma cells could proliferate in response to Dex. In this context, the three-drug combination of a human monoclonal anti-CXCR4 antibody with lenalidomide and Dex or bortezomib and Dex phase Ib/II study demonstrating a high response rate is of particular interest (Ghobrial 2019 phase).

## Materials and Methods

*Molecular Biology*.

### Cell line culture

MM.1S is a multiple myeloma glucocorticoid sensitive cell line (ATCC® CRL-2974^TM^). Cells were cultured in RPMI-1640 supplemented with 10% fetal bovine serum, and 2 mM L-glutamine. Cell line is tested negative for mycoplasma according to the manufacturer’s instructions (PCR Mycoplasma-Test Kit I, ITW Reagent, A9753). Cells were initially cultured for 24 hour in reduced-serum, hormone stripped media (RPMI 1640 Medium, no glutamine, no phenol red, ThermoFisher Scientific, 32404014) with 10% Charcoal/Dextran treated FBS (Charcoal STRP FBS One Shot, ThermoFisher Scientific, A3382101) and 2 mM L-glutamine to a concentration of 1 million cells per mL. Subsequently, Dexamethasone (Dex) (Sigma D4902) was added to the media at 0.1µM for all treatments timepoints and EtOH was used as vehicle control.

### ChIP-seq Procedure

MM.1S cells were exposed to Dex or EtOH for 1 hour and crosslinked with freshly made 1% formaldehyde (ThermoFisher Scientific, 28908) for 15 minutes and quenched with 125 mM Glycine (Sigma-Aldrich, 50046) for 10 minutes. Cells were pelleted and washed in PBS, then pelleted again and stored at −80°C.

ChIP-seq CTCF was performed as previously described (Jin 2018) with the following modifications. Formaldehyde-fixed cells were lysed and chromatin sheared by sonication using a Bioruptor Pico (Diagenode). IP was carried out using the 3µg of polyclonal CTCF antibody (Diagenode, C15410210). DNA from protein-associated complexes and corresponding input samples were washed, eluted and reversed crosslinking by incubation with RNase A (ThermoFisher Scientific, AM2270) and protein digested with Proteinase K (ThermoFisher Scientific, 25530049). Samples were purified with DNA Clean and Concentrator columns (Ozyme, ZD4013) and measured using the Qubit dsDNA HS Kit (ThermoFisher Scientific, Q32851). Libraries were prepared using NEBNext Ultra II DNA Library Prep according to the manufacturer’s instructions ((New England Biolabs, E7103S). Libraries were sequenced using Miseq platform (Kit 150cycles V3-PE) with 20 million reads per sample.

ChIP-seq GR (H-300) (Santa Cruz Biotechnology, sc-8992) and H3K27ac (Active Motif, AM-39133) were performed by Active Motif Epigenetic Services. Sequencing depth was 40 million reads for CHIP-seq GR Dex, 38 million reads for CHIP-seq GR EtOH, 28 million reads for CHIP-seq H3K27ac Dex and 27 million reads for CHIP-seq H3K27ac EtOH.

### RNA-seq Procedure

MM.1S cells were exposed to Dex or EtOH for 4 hours. Total RNA from MM.1S cells was isolated using direct-zol RNA MicroPrep kits (Ozyme, ZR2060) with on-column DNase treatment according to manufacturer’s instructions. Prior to RNA-seq, RNA quality was confirmed on the Agilent Bioanalyzer 2100 using the RNA 6000 Nano Kit (Agilent, 5067-1511). Total RNA-seq libraries were generated using NEBNext Poly(A) mRNA Magnetic Isolation Module (New England Biolabs, E7490S) and NEBNext Ultra II Directional RNA Library Prep (New England Biolabs, E7765S). Libraries were sequenced using the Illumina HiSeq 2500 (Hiseq Rapid SBS kit v2 2*75 cycles).

### Fast-ATAC Procedure

The Fast-ATAC protocol was performed as previously described (Corces 2016) using 0.1 million cells. MM.1S cells were exposed to Dex or EtOH for 1 hour, washed in PBS 1X and centrifuged. The pellet was resuspended in the transposase reaction mix (25µl of 2x TD buffer, 5µl of TDE1, 0.5µl of 1% digitonin, 19.5µl of nuclease-free water) (FC-121-1030, Illumina; G9441, Promega). Transposition reactions were incubated at 37°C for 30 minutes in an Eppendorf ThermoMixer with agitation at 1000 RPM. Transposed DNA was purified using the kit “DNA Clean and Concentrator”-5 (ZD4013, Ozyme). Transposed fragments were amplified and purified as described previously (Buenrostro 2015) with Nextera Index Kit (FC-121-1011, Illumina). qPCR was performed to determine the optimal number of cycles to amplify the library to reduce artifacts associated with saturation PCR of complex libraries. PCR was then performed for the optimum number of cycles using the following PCR conditions: 72°C for 5 min; 98°C for 30 s; and thermocycling at 98°C for 10 s, 63°C for 30 s and 72°C for 1 minute. Libraries were amplified for a total of 11 cycles. Library amplification was followed by solid phase reversible immobilization methodology (SPRI) size selection to exclude fragments larger than 1,200 bp. Libraries were sequenced using Illumina HiSeq 2500 (Rapid Run HiSeq paired-end 2*75cycles).

### HiChIP Procedure

MM.1S cells were exposed to Dex or EtOH for 1 hour, were pelleted and resuspended in freshly made 1% formaldehyde (ThermoFisher Scientific, 28908) at a volume of 1 mL of formaldehyde for every one million cells. Cells were incubated at room temperature for 10 minutes with rotation. Glycine (Sigma-Aldrich, 50046) was then added to a final concentration of 125 mM to quench the formaldehyde. Cells were incubated at room temperature for 5 minutes with rotation. Cells were pelleted and washed in PBS, then pelleted again and stored at −80°C.

The HiChIP protocol was performed as previously described (Mumbach 2016) using 7.5µg antibody to H3K27ac (Diagenode, C15410196) with the following modifications. Samples were sheared using Bioruptor Pico (Diagenode), the amount of Tn5 (Illumina, 15027865) used and number of PCR cycles performed were based on the post-ChIP Qubit amounts. Libraries were sequenced on NovaSeq 6000 (NovaSeq 6000 S1 Reagent Kit 2*100 cycles).

### Single Cell RNA-seq Procedure

For scRNA-seq, MM.1S cells were exposed to Dex or EtOH for 4 hours and 24 hours. Single cell RNA-seq profiling was performed with the ChromiumTM Single Cell Controller. A total of 6,000 cells were loaded per lane and processed for complementary DNA synthesis and library preparation, per the manufacturer’s protocol using 3’ v3.1 chemistry (10X Genomics – 1000121). Libraries were sequenced on NovaSeq 6000 (NovaSeq 6000 S1 Reagent Kit 2*100 cycles) to a mean depth of 45,000 reads/cell using the read lengths 26bp Read1, 8bp i7 Index, 98bp Read2.

### Single Cell Multiome ATAC + Gene Expression Procedure

For scMultiome, MM.1S cells were exposed to Dex or EtOH for 1 hour and 4 hours. Single cell 3’ gene expression and open chromatin libraries were simultaneously generated using Chromium Next GEM Single Cell Multiome ATAC + Gene Expression Kit from 10x Genomics, following the protocol provided by the manufacturer. A total of 5,000 nuclei were loaded per lane on the ChromiumTM Single Cell Controller. Libraries were sequenced on NovaSeq 6000 (NovaSeq 6000 SP Reagent Kit 2*50 cycles) to a minimum depth of 24,000 reads/nucleus for Gene Expression library and 42,000 reads/nucleus for ATAC library.

### Rapid immunoprecipitation mass spectrometry of endogenous protein (RIME)

RIME GR (H-300) (Santa Cruz Biotechnology, sc-8992) was performed by Active Motif Epigenetic Services. MM.1S cells were exposed to Dex or EtOH for 1 hour and fixed according to the manufacturer’s instructions (RIME Cell Fixation protocol, Active Motif). Analysis were performed by Active motif and resuslts are given as supplementary table (see Additional file 2: Table S1).

### Computational analysis

#### Chromatin state annotation

To obtain functional annotation of MM.1S cell line, we used ChromHMM (v1.11) (Ernst 2010; Ernst 2012). Five histone marks available from ENCODE consortium (Consortium 2012) (H3K4me1, H3K4me3, H3K27ac, H3K36me3 and H3K27me3) in three different cell lines (MM.1S, U266 and GM12878) were analyzed using hidden Markov model to identify 10 different chromatin states. Default parameters of chromHMM were used. Bam files were binarized into 200bp genomic windows and the presence or absence of each histone mark was evaluated. Then we employed biological analysis to annotate those chromatin states giving them biological meanings.

#### Treatment of ChIP-seq data

ChIP-seq sequencing quality was assessed with fastqc (v0.11.8) (Andrews 2010). ChIP-seq read adaptors were firstly trimmed using trimmomatic (v0.39) (Bolger 2014) and then reads were mapped using bowtie2 (v2.1.0) (Langmead 2009) to the Human genome UCSC hg19 (GRCh37) (Kent 2002). Only one mismatch was allowed. After alignment step, unmapped reads, low quality mapped reads (mapQ<30) and reads mapped to ENCODE blacklist regions (Amemiya 2019) were removed with samtools (v1.3.1) (Li 2009) for analysis. We also removed reads that were like to be optical and/or PCR duplicates using picard MarkDuplicates (v2.23.5) from GATK (Mckenna 2010).

ChIP-seq enriched regions defined as peaks were called using macs2 (v2.1.1) (Zhang 2008) versus input (sequencing without immunoprecipitation). We only retained peaks higher than specified p-value threshold (pval<1e^-07^).

#### Treatment of ATAC-seq data

All ATAC-seq data were processed based on Kundaje lab proposed pipelines (Koh 2016; Liu 2019) available on github. Quality of sequencing assessment, read adaptors trimming, read mapping to Human genome hg19 and filtration were performed the same way as ChIP-seq reads. Before peak calling steps, and due to the Tn5 insertion, mapped reads were shifted with respectively 5bp and 4bp for strand + and strand - with samtools. Finally, enriched regions defined as ATAC-seq peaks were called using macs2 only significant peaks were retained (FDR<0.05).

#### RNA-seq differential analysis

Each RNA-seq sample was mapped using Tophat2 (Trapnell 2009) versus hg19 reference genome. We then employed proposed protocol (Trapnell 2012) to perform differential expression analysis with cufflinks. Only genes with a LogFC greater than or equal to 0.6 and an FDR<0.05 were kept for analysis.

#### HiChIP data treatment and differential analysis

We employed HiC-pro (Servant 2015) to process HiChIP data from raw-data to normalized contact maps. All reads were mapped to hg19 genome using bowtie2 (global parameters: --very-sensitive -L 30 --score-min L,-0.6,-0.2 --end-to-end -- reorder; local parameters: --very-sensitive -L 20 --score-min L,-0.6,-0.2 --end-to-end --reorder). Contact maps were generated at different resolution (1kb, 2kb, 15kb, 20kb and 40kb) and normalized by the iterative correction and eigenvector decomposition (ICED) method. HiC-pro output directory was then used as input to Hichipper (Aryee) with MboI restriction site position for loop calling.

Differential analysis of chromatin loops was performed with function exactTest of package edgeR (Robinson 2010), with default parameters except for dispersion, which was set to “trended”. Interaction with FDR below 5% and absolute logFC above 0.60 were considered significant.

#### Global treatment of genomic data

Genomic data were proceeded using different genomic tools such as Bedtools (v2.28.0) (Quinlan 2010) for manipulating genomic files, the homer suite for annotation and motif discovery (v4.4) (Heinz 2010). Data were also treated using own Python scripts (v2.7).

#### Motif search

De novo motif discovery was performed using the MEME suite (v4.11.2) (Bailey 2015) for GR peaks centralized on peak submit and extend with 250bp in both directions. Motif from 6bp to 16bp were searched with a maximum of 5 motifs were asked. To identify sequences where a specific motif is found, we employed FIMO tool from MEME. Finally, to identify centrally enriched motif, we used centriMo from MEME.

#### Signal tracks generation

We employed the bamCoverage tool from the Deeptools (v2.0) (Ramirez 2014) suite to generate bigWig files. Signal tracks files were normalized using RPGC method (Read Per Genomic Content) also known as the 1X normalization included in bamCovergae options.

Once those files were generated, we used the bigwigCompare tool to create a differential tracks between H3K27ac with or without Dex.

All ChIP-seq and ATAC-seq files were generated with this method. Visualization of signal tracks were obtained using the Integrative Genome Viewer IGV (Robinson 2011).

#### Genome ontology analysis

Genome ontology analysis was performed using GREAT (v3.0.0) (McLean 2010) with default parameters (Gene regulatory domain: prox. 5kb upstream and 1kb downstream; dist. up to 1000kb). Enrichment statistics were computed using binomial and hypergeometric gene-based test. Pathways were selected as significantly enriched if the false discovery rate (FDR q-value) was lower than 0.01.

#### Differential analysis of ChIP-seq H3K27ac peaks

In order to find H3K27ac ChIP-seq responding to GR binding, we first selected all H3K27ac peaks found within GR peaks (n=16,228). On those sites, we then estimated the normalized count (RPGC: Read Per Genomic Content) of H3K27ac ChIP-seq in both conditions. Log2-Fold changes were then calculated for each site and we consider as H3K27ac Dex-increased all sites with a log2FC higher than 0.1. The H3K27ac Dex-increased peaks are given as supplementary table (see additional file 3 - Table S2).

#### Identification of the GR-predominant enhancer among each regulatory network of Dex-responsive genes

We first collected all the H3K27ac Dex-increased distal-proximal loops linked with an H3K27ac Dex-increased ChIP-seq peaks. Overlapping with proximal anchors of Dex-induced genes highlights a subset of 55 Dex-responsive genes. To find the predominant enhancer in each regulatory network of those 55 Dex-responsive genes, we proceeded in three steps. First, we selected the anchor loop closest to the TSS of each up-regulated gene linked with increased chromatin loops. Then, for each gene, we collected all anchors linked to a loop to this specific TSS anchor. Finally, for each gene, we selected among the regulatory network the anchor with the highest GR ChIP-seq signal referred as the predominant enhancer while the other ChIP-seq peaks found within regulatory network were considered as the supportive enhancers. All 55 Dex-responsives genes GR predominant enhancer are given as supplementary table (see additional file 5 - Table S4).

#### Differential analysis of ATAC-seq peaks found within chromatin loop anchors

We collected all ATAC-seq peaks found within chromatin loops anchors and, for each peak, we estimated the RPGC count of ATAC-seq in EtOH and Dex conditions. Log2FC were then estimated and all ATAC-seq peaks with a LogFC greater than or equal to 0.6 were considered as ATAC up. The ATAC Dex-increased peaks obtained are given as supplementary table (see additional file 6 - Table S5).

#### Global treatment of single cell data

Preprocessing steps for single cell data were done using CellRanger Software suite, respectively cellranger (v5.0.0) (Zheng 2017) and cellranger-arc (v1.0.1) for scRNA-seq and scMultiome-seq (Satpathy 2019). For both type of data the hg38 genome assembly provided by 10xGenomics was used for alignment. Further analyses were performed on R (v3.6). For scRNAseq, Count matrices were loaded into R using the Seurat package (v3.9.9) (Satija 2015). For each cell we calculated the percentage of mitochondrial reads (percent.mt) and the percentage of nuclear retained lncRNA (percent.nc). We also used the CellCycleScoring function from Seurat to assign a cell cycle state to each cell (Phase), the assignement of the cell cycle state is based on the S.score and G2M.score calculated by this function. Cells were then filtered on the following criteria: 5<percent.mt<25, percent.nc<10, a minimum of 2,000 reads and 1,500 different genes expressed. Normalization and dimensional reduction were performed using the Seurat NormalizeData function with standard parameters. The function FindVariableFeatures was then used to select the 3,000 most variable features, those features have been scaled with Seurat ScaleData function, because cell cycle was a major part of the variability, we added S.score and G2M.score to the vars.to.regress argument of the function. We reduced dimension using RunPCA from Seurat, only the first 30 dimensions were used for downstream analyses. We also calculated a 2D embedding of our cells with RunUMAP, neighbor search and clustering were performed using FindNeighbors and FindClusters functions, with default parameters. For scMultiome-seq, RNA and ATAC matrices were loaded into R using Seurat and Signac (v1.1.0) (Stuart 2020) packages. For each cell we calculated the percent.mt, percent.nc and the Phase. For the ATAC data we also calculated transcription starting site (TSS) score and the ratio of reads overlapping with the blacklisted regions of the genome contained in the blacklist hg38 unified provided by the Signac package. Cells detected by cellranger were filtered on both RNA and ATAC data. For RNA data, we kept cells between 3,800 and 150,000 reads and more than 2,000 different genes expressed. We also kept cells with a percent.mt between 5 and 30 and a percent.nc lower than 8. For ATAC data, we kept cells with a number of reads between 10,000 and 500,000 and a number of different features between 5,000 and 60,000. We also filtered cells with a TSS enrichment between 3.5 and 15, a nucleosome signal lower than 1,5 and more than 50% percent reads in peaks.

For normalization and dimensionality reduction we used the RunTFIDF and RunSVD functions from the Signac packages. RunSVD was run on the features selected by FindTopFeatures with min.cutoff set to q80. UMAP embedding, neighbor search and clustering were performed the same way as for the RNA data alone.

Differential expression and accessibility were tested using the findMarkers function provided by Seurat, with test.use argument respectively set to “MAST” and “LR”.

#### Assesment of scATA-seq peaks co-accessibility

Co-accessibility scores between scATACseq peaks were calculated using Cicero (v1.3.4.11) (Pliner 2018) with default parameters. Co-accessibility tables were built on each treatment condition separately. From those tables of co-accessibility scores we built two networks for each of the 55 Dex-responsive genes GR-predominant enhancers on one hand and GR-supportive enhancer in other hand. Nodes of those networks were defined as all peaks overlapping with the selected regions. Edges were built using the co-accessibility tables, considering only connections with a score higher than 0.1.

#### Single cell Dex-activated Genes (scDAGs)

We studied the distribution of logFC above 0 and found out it was bimodal with a small part of positive logFC being far from the main part. We then used gaussian mixture model to identify the small sub-population of high logFC genes, i.e. scDAGS.

#### Ratio of responding genes

For each gene, we computed the 80e percentile of expressed values in untreated cells; we then calculated, for each cell, the Ratio of Responding Genes (RRG) as the percentage of scDAGs with expression value above the gene threshold for untreated cells, 4 hours Dex treated cells and 24 hours Dex treated cells.

## Supporting information

Additonal File 5 - Table S4

Additonal File 4 - Table S3

Additonal File 3 - Table S2

Additonal File 2 - Table S1

Additonal File 1 - Figure S1-S15

## Acknowledgements

We thank the Genomics and Bioinformatics core facility of Nantes (GenoBiRD, Biogenouest, IFB) for its technical support.

## Abbreviations

GC: Glucocorticoids

GR: Glucocorticoid Receptor

GRE: Glucocorticoid response element

MM: Multiple Myeloma

Dex: Dexamethasone

IMiDs: immunomodulatory drugs

PIs: Proteasome Inhibitors

TF: Transcription Factor

SE: Super-Enhancers

ISRE: Interferon Signaling Response Element

scDAGs: single cell Dex-Activated Genes

RRG: Ratio of Responsive Genes

TSS: Transcription Start Site

kb: kilobases

IQR: Inter-Quartile Range

FPC: Fraction of Positive Cells

FDR: False Discovery Rate LogFC:Log Fold Change.

## Availability of data and materials

ChIP-seq, HiChIP-seq, ATAC-seq, RNA-seq, scRNAseq and scMultiome have been deposited at the European Genome-phenome Archive (EGA, https://www.ebi.ac.uk/ega), wich is hosted bye the EBI and the CRG, under dataset accession EGAD00001007927 (https://ega-archive.org/datasets/EGAD00001007927), EGAD00001007926 (https://ega-archive.org/datasets/EGAD00001007926), EGAD00001007925 (https://ega-archive.org/datasets/EGAD00001007925), EGAD00001007928 (https://ega-archive.org/datasets/EGAD00001007928), EGAD00001007929 (https://ega-archive.org/datasets/EGAD00001007929) and EGAD00001007924 (https://ega-archive.org/datasets/EGAD00001007924), respectively.

## Ethics approval and consent to participate

Not applicable.

## Consent for publication

Not applicable.

## Authors’ contributions

S.M. and F.M. designed the study, collected, and analyzed the data and wrote the paper; V.G., J.C., J.B. and C.H. collected and analyzed the data and wrote the paper; JB.A., E.D., N.R., M.D., F.W. and P.M. collected the data; C.G.-C. and L.C. analyzed the data.

## Additional Files

Additional file 1 --- Figure S1-S15 Supplementary figures.

Additional file 2 --- Table S1

Enriched Proteins List for GR RIME Final Assay Additional file 3 --- Table S2

H3K27ac Dex-induced enhancers Additional file 4 --- Table S3

41 Dex-responsive genes with increased proximal anchor and Dex SE Additional file 5 --- Table S4

Predominant enhancer associated with its target Dex-responsive gene Additional file 6 --- Table S5 ATAC-seq peaks Dex-increased linked to increasing chromatin loops

